# *Hyobanche hanekomii* (Orobanchaceae) is pollinated by non-flying mammals and birds

**DOI:** 10.1101/2022.12.13.519978

**Authors:** Tim Niedzwetzki-Taubert

## Abstract

*H. hanekomii* was recently described as a new member of the genus *Hyobanche* inside the Orobanchaceae family. *H. hanekomii* is a small geoflorous holoparasitic plant, which often grows under shrubs and has red-flowered inflorescences, which do not emit a scent. The plant combines characteristics from bird- and mammal-pollinated plants. Until now it was uncertain if *H. hanekomii* is pollinated by birds or by non-flying mammals, since the species is showing an intermediate morphology when compared to *H. atropurpurea* (mammal pollinated) and *H. sanguinea* (bird pollinated)*. Elephantulus edwardii* and *Nectarinia famosa* were observed foraging on *H. hanekomii* inflorescences indicating a mixed pollination syndrome or a transmission state between bird pollination to pollination by non-flying mammal. In this study I prove that *E. edwardii* and *N. famosa* are indeed pollinators of *H. hanekomii*. This was done by observing the interaction between different animal species and *H. hanekomii* inflorescences in their natural habitat by using camera traps and camcorders.

It could be observed that *E. edwardii* licked nectar from flowers of *H. hanekomii* with its long tongue. The animals pressed their rostra deep into the flowers. While foraging on the flowers, it could be observed that pollen was placed onto the rostra of the animals. It was also observed, that the stigma of the flower touched the animals on the same spots where pollen was placed. To drink on the flowers, *Nectarinia famosa* inserted its beak into the flowers. While doing so, it was observed that pollen was placed on the beaks and that the stigma touched the animals. It is possible that both species transport pollen from one *H. hanekomii* flower to another while foraging on them, and that the pollen reaches the stigma of another flower. Because of this both species are considered as pollinators of *H. hanekomii*.

*E. edwardii* was identified as the main pollinator of *H. hanekomii* as they visit the inflorescences frequently. *N. famosa* was detected as a secondary pollinator of *H. hanekomii* as they visit the flowers infrequently. This issue has to be examined further since the rarity of *N. famosa* visits could be influenced by different factors like removal of perching positions around the plants to have a better camera angel or by the cameras itself, so that *N. famosa* would visit the *H. hanekomii* inflorescences more often in an non altered surrounding.

In addition UV photography and spectrometry revealed that no UV reflecting areas are present on the plants indicating adaption to non-flying mammal than bird pollination syndrome. The same counts for missing stinging hairs on the flowers. In contrary some of the important flower characteristics (Flower entrance width and height) are significant smaller than those of the non-flying mammal pollinated plant *H. atropurpurea* indicating a bird pollination syndrome.

## Introduction

Pollination is the transfer of pollen from an anther to a stigma of a flowering plant, whereas pollination by wind, water, and animals (Zoophily) are the most important forms and 75 percent of all angiosperms have specialised on the later (Faegri & Van der Pijl, 1979; Raven, P.H., Evert, R.F., 2006; Sadava, D., Hillis, D.M., Heller, H.C., Berenbaum, M.R., Markl, 2011). Zoophily is divided into ornithophily, entomophily, chiropterophily, and pollination by non-flying mammals (Raven, P.H., Evert, R.F., 2006; Sadava, D., Hillis, D.M., Heller, H.C., Berenbaum, M.R., Markl, 2011; Townsend, C.R., Begon, M., Harper, 2014). The pollination of a plant species by certain animal groups results in adaption of a plant to this group, which is described as pollination syndromes (Fenster *et al.*, 2004). The non-flying mammal pollination syndrome covers primates, rodents, marsupial and other small mammals (Carthew and Goldingay, 1997; Prance, 1980; Steenhuisen *et al.*, 2015; Wester, 2011, 2015; Wester *et al.*, 2009; Yumoto, 2000; Zoeller *et al.*, 2016).

Pollination syndromes describe flower characteristics like morphology, colour, nectar, and scent which attract distinct pollinators, allowing them to forage on the flowers, while excluding illegitimate flower visitors from foraging as they do not contribute to the pollination of the species (Fenster *et al.*, 2004; Ollerton, 1998).

Plants which are specialised on the pollination by non-flying mammals express properties such as the absence of stinging or burning –hairs, geoflory, open, bowl shaped, stable flowers with short stalks, stable flower organs, a lot of low concentrated nectar, a lot of pollen, flowers with a big distance between stigma and nectar, special scent which is often nutty, buttery, or yeasty, inconspicuous green-brown to dark coloration, clustered flowers and a flowering time where other recourses are scarce (winter) as well as cryptic inflorescences (Biccard and Midgley, 2009; Cocucci and Sérsic, 1998; Melidonis and Peter, 2015; Rourke and Wiens, 1977; Wiens *et al.*, 1983). Additional indicators for pollination by non-flying mammals are non-destructive feeding on flowers, evidence of pollen on animals or in their faeces, contact between pollen which is deposited on the animal and the stigma of a flower, only infrequent flowers visits by other animal groups, reduced fertility if non-flying mammals are restrained from the plant as well as the observation, that nectar, scent production and flowering time are consistent with the periods of pollinator activity (Johnson *et al.*, 2011; Rourke and Wiens, 1977, 1978; Wiens *et al.*, 1983). This pollination syndrome was observed all over the world, for example in Australia, New Guinea, Malaysia, China, India, Madagascar, South Africa, Brasilia, Peru, and Costa Rica (Carthew and Goldingay, 1997; Collins and Rebelo, 1987; Goldingay *et al.*, 1991; Janson *et al.*, 1981; Kress *et al.*, 1994; Lumer, 1980; Lumer and Schoer, 1986; Prance, 1980; Wester *et al.*, 2009; Wiens *et al.*, 1979; Yumoto, 2000). Around 100 plant species are known which are visited by more than 50 different non-flying mammals and recently even lizards were observed play a role in pollination (Carthew and Goldingay, 1997; Johnson *et al.*, 2001; Wester, 2019). Plants which are specialised on the pollination by birds in contrast show properties such as red, yellow, orange or similar bright coloured flowers, have lots of thin liquid nectar, emit no scent, have long flower tubes or nectar spurs, and small flower openings as well as perching positions, are good visible and robust, and have their anthesis in daytime (Cronk and Ojeda, 2008; Faegri & Van der Pijl, 1979; Johnson and Nicolson, 2008; van der Pijl, 1961, 1969; Raven, P.H., Evert, R.F., 2006; Strelin *et al.*, 2016).

Pollination syndromes can be used to make assumptions about which group of animals can pollinate a plant, however it should be viewed critically and checked by field observations or experiments as the classification into certain syndromes is often incorrect (Fenster *et al.*, 2004; Ollerton, 1998; Ollerton *et al.*, 2009).

Pollination by non-flying mammals is one of the least researched pollinator phenomena. In South Africa this pollination syndrome was firstly observed in 1978 and is known for species from the Ericaceae, Proteaceae, Hyacinthaceae, Colchicaceae, Fabaceae, Cytinaceae, Xanthorrhoeaceae und Asparagaceae (Biccard and Midgley, 2009; Collins and Rebelo, 1987; Johnson *et al.*, 2001; Kleizen *et al.*, 2008; Letten and Midgley, 2009; Payne *et al.*, 2016; Rourke and Wiens, 1978; Turner *et al.*, 2011; Wester, 2015; Wester *et al.*, 2009). Within those plant families different rodent species like *Acomys subspinosus, Aethomys namaquensis, Myomyscus verreauxii, Rhabdomys pumilio, Gerbillurus paeba*, and *Mus minutoides* are identified as the pollinators (Biccard and Midgley, 2009; Johnson *et al.*, 2001; Letten and Midgley, 2009; Rourke and Wiens, 1978; Turner *et al.*, 2011; Wester, 2015; Wester *et al.*, 2009). But other non-flying mammals like elephant shrews *Elephantulus edwardii*, *Elephantulus brachyrhynchus*, *Elephantulus myurus* (Macroscelidea) can serve as pollinators too, which was just proven recently under laboratory conditions and in the wild for *Hyobanche atropurpurea* (Orobanchaceae), *Whiteheadia bifolia, Massonia echinata* (Asparagaceae), *Aloe peglerae* (Xanthorrhoeaceae), and *Cytinus visseri* (Cytinaceae) (Flasch *et al.*, 2017; Johnson *et al.*, 2011; Payne *et al.*, 2016; Wester, 2010, 2011, 2015).

Ornithophily has evolved several times over the course of evolutionary history and is known in more than 50 bird families, which pollinate flowers from 65 plant families worldwide by feeding on the nectar of the flowers and thus contribute to their pollination (Cronk and Ojeda, 2008; Geerts and Pauw, 2009; de Waal *et al.*, 2012). In South Africa ornithophily is known for plant species from the Apocynaceae, Xanthorrhoeaceae, Orobanchaceae, Hyacinthaceae, Cytinaceae, Orchidaceae, Penaeaceae, Retziaceae, Iridaceae, Lamiaceae und Proteaceae which are pollinated by hummingbirds (Trochilidae), sunbirds (Nectariniidae), and honeyeaters (Meliphagidae) (Collins and Rebelo, 1987; Geerts and Pauw, 2009; Hobbhahn and Johnson, 2015; Johnson, 1996; Johnson and Brown, 2004; Pauw, 1998; Payne *et al.*, 2016; Steenhuisen *et al.*, 2012; Turner and Midgley, 2016; de Waal *et al.*, 2012; Wester and Claßen-Bockhoff, 2006). Various species of sunbirds can be considered as pollinators of a plant species, while plants in the South African region have been observed to exhibit adaptions to either long or short beaked sunbird species as nectar robbing by short beaked sunbird species have been observed (Geerts and Pauw, 2009; de Waal *et al.*, 2012).

The genus Hyobanche is endemic to southern Africa with only 10 described species (Wester, 2011; Wolfe, 2018, 2013; Wolfe and Randle, 2001). As holoparasites the plants take up nutrients from their host plants via haustoria and have large underground rhizome networks, while only the inflorescences of the plants grow above the ground and are often hidden under other plants (Morawetz *et al.*, 2010; Wester, 2011; Wolfe, 2013; Wolfe and Randle, 2001). *H. atropurpurea* is known to be pollinated by *E. edwardii*, has dark almost black flowers with wide antechambers and emits a musty odour, while *H. sanguinea* on the other hand is pollinated by sunbirds, has bright showy red flowers with narrow inflorescences and does not emit any fragrance (Turner and Midgley, 2016; Wester, 2011). The recently described species *H. hanekomii* (Wolfe, 2018) - which is endemic to the Cape Fold Mountains of the Western Cape (between Citrusdal and Van Rhynsdorp) and grows in the Sandstone fynbos biome-shows an intermediate morphology compared to the other two species and could potentially be pollinated by either birds, small non-flying mammals or both (Mucina and Rutherford, 2006; Wolfe, 2018). As the species combines different traits from both avian and mammalian pollination syndromes further investigation is necessary to determine the pollinator. Based on the red flower colour (Figure 1, A and B) and the lack of fragrance (personal experience), it can be hypothesized that *H. hanekomii* is bird pollinated. However, the size of the flower entrances, the geoflory and crypsis of the plants (Figure 1, A to E) and the growth which is typical for *H. atropurpurea* (Figure 1, C) do not exclude mammals as possible pollinators. The flower colour of *H. hanekomii* is quiet variable (compare Figure 1, A, B, D, E) from light red-purple to very dark, almost black red, giving this species a colour spectrum between the bright *H. sanguinea* and the almost black *H. atropurpurea*.

**Figure 1:**
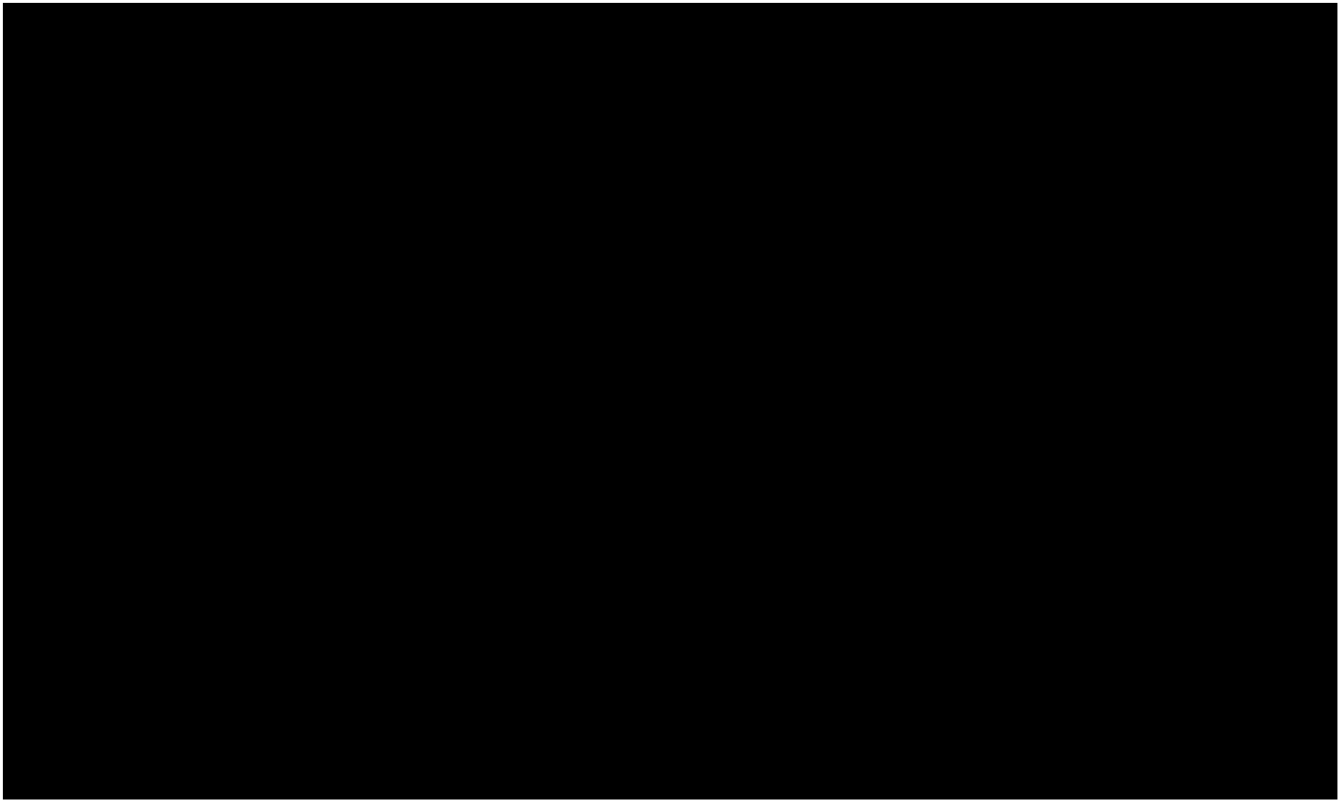
Hyobanche Inflorescences and flowers. **A & B:** *H. hanekomii* inflorescences with differential coloration. **C:** *H. atropurpurea* inflorescence. **D & E:** Single flowers of *H. hanekomii* with great variation in the size of the flower entrance and coloration. Pictures taken by Niedzwetzki-Taubert, T., Karvang, A.S.N., Wester, P. **Pictures not shown due to copyright reasons**

Based on this discrepancy this work deals with the question of whether *H. hanekomii* is pollinated by non-flying mammals, by sunbirds, or even by both. To answer this question *H. hanekomii* inflorescences were observed using wildlife cameras and camcorder to evaluate the footage for potential pollen transfer and the results were discussed in the context of previous findings. The data was evaluated with regard to the feeding behaviour of different animal species visiting the flowers, the contact of the animals with the reproductive structures of the plants and potential pollen transfer. Additionally flower traits were analysed to see if they correlate more with bird or mammal pollinated species of the genus.

## Material and methods

The video observations used to identify the flower visitors of *H. hanekomii* and their behaviour were recorded on the premises of the Fonteintjie Farm (S32°37.237’ E019°03.228’ approximately 6.5 kilometre southeast of Citrusdal in the Cederberg Mountains, Western Cape, South Africa) over a period of 16 days from 08.09.2016 to 23.09.2016 in an area of 16.875 m^2^.

Observations were made using up to 25 wildlife cameras (Nature View Cam HD MAX 119440, Bushnell, Cody, USA). Records were made during day and night (using the infrared sensor of the wildlife cameras). Close-up lenses were used, which enabled working distances of 0.3 m, 0.5 m, 1.2 m, and 2.5 m to the plants. The trigger delay was set to 0.7 seconds after triggering caused by motion and heat. Each shot was 60 seconds long, and there was a one second delay between shots. Additionally six video cameras (HDR-XR520 and HDR-XR550, Sony, Tokyo, Japan) were set up, which recorded up to 13 hours straight. The total running time was 7875 hours, were 60 hours were recorded by video cameras at daylight and 7816 hours were recorded by wildlife cameras with roughly a 1:1 ratio of day and night recordings. In total 67 different *H. hanekomii* inflorescences were observed, while between one and five inflorescences were filmed per camera.

All videos were viewed using DivX Pro Media Player 10.7.1 (DivX, LLC, San Diego, USA). Videos in which animal activity was detected were then evaluated in detail on the foraging bout by using the Free Video Editor 1.4.53 (DVDVideoSoft, London, England). A foraging bout was defined as the period from the beginning to the end of an interaction between an animal and a plant(s). If a camera was triggered several times in a row and it was clearly the same animal, this was rated as a single foraging bout. During a foraging bout it was analysed with how many inflorescences and flowers an animal interacted, how many flowers an animal drank or sniffed on, and if possible the frequency with which the animal licked up the nectar was determined. Furthermore the spatial relationship between the nectar drinking animal and the flowers and flower organs was analysed, and how long an animal occupied itself with the plants and flowers. It was also analysed whether pollen was transferred to animals and whether there was contact between the animals and the stigmas of the flowers.

The different animal species were identified using fauna and flora in Southern Africa (Apps *et al.*, 2001) and bird forums (http://www.birdforum.net/opus/Malachite_Sunbird, 30.12.2016; http://gerdandre.de.tl/Singv.oe.gel.htm, 30.12.2016). Different mouse species could not be distinguished in the nocturnal video recordings, since the best point of differentiation is the coat colour, which was not recognizable.

Statistical analysis was carried out with “R” 3.3.2 (31.10.2016) (R Development Core Team, 2016). The data sets were tested for normal distribution using Shapiro-Wilk tests. If two data sets were compared with one another, a t-test was carried out for normally distributed data sets and a Mann-Whitney U-test (Wilcox test) for non-normally distributed data sets. If more than two sets of data were compared, an ANOVA was performed for normally distributed data and a Kruskal-Wallis test for non-normally distributed data. A pairwise Wilcox test was then performed, in which the p-value was adjusted using the False Discovery Rate method to minimise type 1 error.

To access the presence of ultraviolet light reflecting pattern on the plants UV photography was used. For this approach a modified Nikon D40 digital camera (Nikon, Tokyo, Japan) was used in which the low pass filter was replaced by an UVI filter (Modified by Optik Makario, Mönchengladbach, Germany). As an objective the Nikkor 18-55mm f/3.5-5.6G VR (Nikon, Tokyo, Japan) was used. Different filter were used in combination with the camera. For UV photography the UV400N filter and for normal pictures a IR-neutralisation filter NG was used. Additionally a UV lamp (Nite Hunter, UV 365nm Wolf-eyes, Shenzhen, China) was used to illuminate plants and flowers to ensure enough UV light for reflection.

To access the colour spectrum of H. hanekomii the mobile JAZ USB Spectrometer (Ocean Optics, Dunedin, USA) was used to analyse 1 flower from 32 different plants. The average and standard deviation of the spectra was calculated.

To compare the morphology different plant and flower traits of 31 *H. hanekomii* inflorescences and 11 *H. atropurpurea* inflorescences were measured with a sliding calliper and compared to each other. A two sided heteroscedastic t-test was used to evaluate significant differences between the two species.

## Results

Using the video files elephant shrews (*E. edwardii*), malachite sunbirds (*N. famosa*) and different mice species (Muridae) could be identified interacting with *H. hanekomii* inflorescences. Beaked turtles (Chersina angulata), chacma baboons (Papio ursinus), and domestic cattle (Bos primigenius taurus) were observed destroying and eating whole plants or parts of the plant, and can therefore be excluded as potential pollinators.

*E. edwardii* could be observed between 18:16 and 07:58, as a regular flower visitor on *H. hanekomii* inflorescences on all 16 observation days. If *E. edwardii* noticed a *H. hanekomii* inflorescence the animal ran towards it, sniffed at the flowers or drank their nectar. Here the animals entered the flower openings with their rostrums and licked the nectar out of the flower chambers with their long tongues (Figure 2, A).

**Figure 2:**
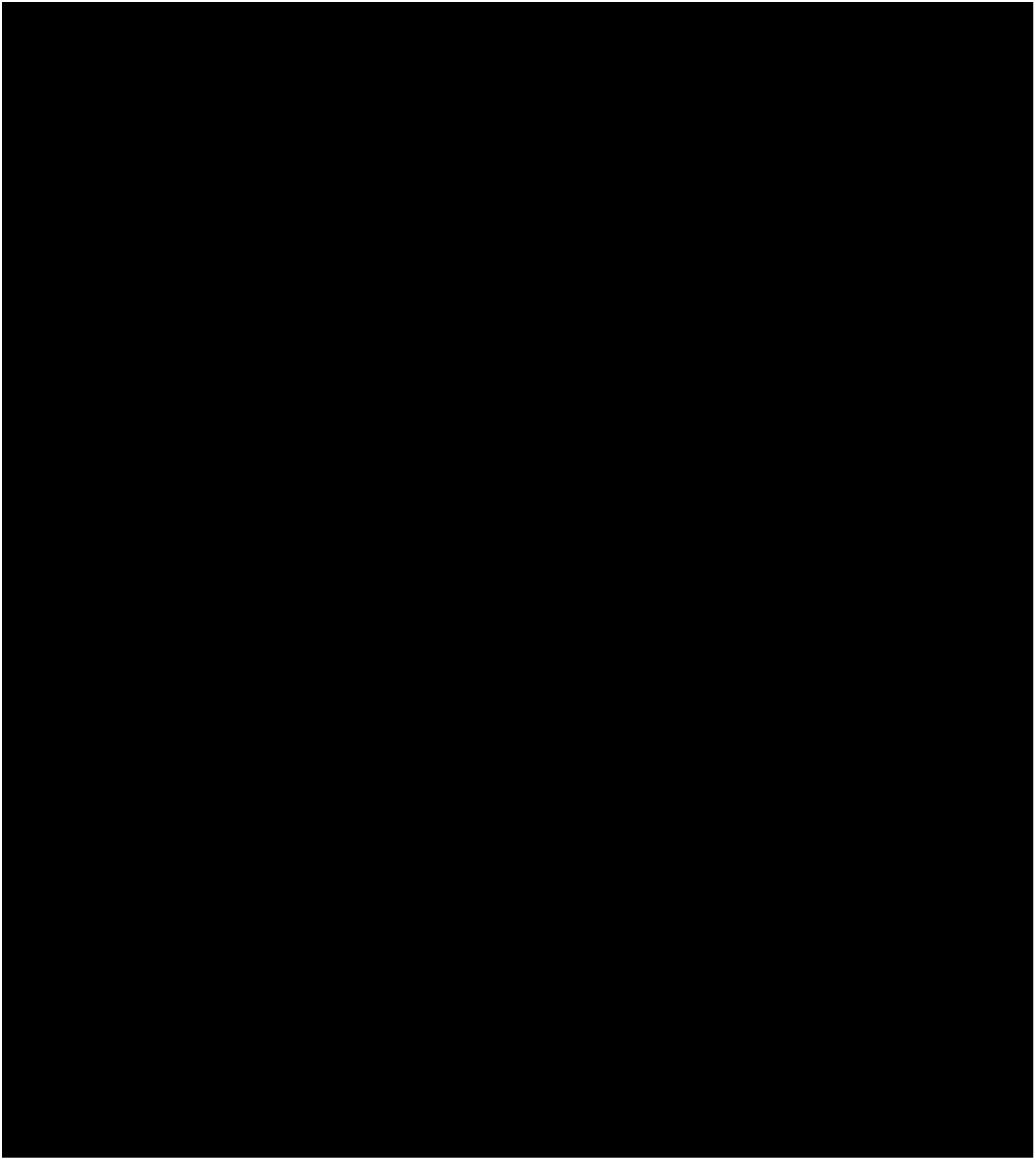
*E. edwardii, N. famosa*, and a rodent visiting *H. hanekomii* inflorescences. **A:** *E. edwardii* drinks from a flower, with the stigma of the flower touching the animal’s rostrum and forehead. **B:** *E. edwardii* after visiting a flower with pollen on the rostrum, which is recognisable as a white spot. **C:** *N. famosa* is drinking nectar from a flower. **D:** *N. famosa* after a flower visit with clearly visible pollen on the beak attachment point. **E:** A rodent is sniffing on a flower. During this process the animal brings its head into close proximity to the flower entrance (stigma and pollen sacks above the head) while nearly pressing the head into the flower. Pictures taken by Niedzwetzki-Taubert, T., Karvang, A.S.N., Wester, P. **Pictures not shown due to copyright reasons**

A total of 10331 flower visits by *E. edwardii* were observed where nectar was drunk. Additionally 2130 flower visits were observed at which *E. edwardii* individuals only sniffed. When drinking from a flower the animal stood crouched from the front to the side (90° angle) with its head tilted in front of the flower. When *E. edwardii* drank from the flowers head-on pollen could be deposited all over the animal’s proboscis and all over the snout up to below the eyes (Figure 2, B). It was observed, that the stigma touched the animals in the same areas. If *E. edwardii* drank from the side of the flowers pollen was deposited on the proboscis as well as on its sides and on the cheeks of the animal up to the eyes. Stigma touches were also possible in these areas. Destruction of flower organs by *E. edwardii* could not be detected at any flower visit. *E. edwardii* also did not eat pollen from the plants or used its paws as a tool to enlarge the flower entrances.

When a flower had nectar and *E. edwardii* drank from it the flower visit lasted an average of 4.5s (±2.9s). *E. edwardii* tested whether a flower contained nectar or not by sniffing or by inserting the tongue into the flower once, whereby pollen could be transferred to the animals or the stigma touched the animals which was observed rarely ~ 10 times. The vigorous licking of the animals caused the inflorescences to wobble. As a result pollen from the anthers was released onto the animal. The frequency with which *E. edwardii* took up nectar averaged 4.3Hz (±1.7Hz). 548 inflorescence visit by *E. edwardii* were observed, with an average visit of 15.6s (±14.5s). *E. edwardii* drank only once from a flower during an inflorescence visit (Wilcox-Test, p=0,711), but sniffed on a flower more than once per inflorescence visit (Wilcox-Test, p<0.001). A foraging bout of *E. edwardii* was with 38.2s (±27.1s) significant longer than an inflorescence visit of the species lasted on average (Wilcox-Test, p<0.001), which lead to the conclusion that *E. edwardii* visited several inflorescences and often changed them during a foraging bout, but often came back to the plants. For *E. edwardii* 244 foraging bouts and 53 sightings in inflorescence proximity without interactions were documented, which lead to an interaction rate of 82.2 %. Per foraging bout an average of 4.2 (±3.3) flower visits took place from which nectar was drunk. Per foraging bout an average of 8.7 (±6.3) flower visits were made to sniff. This led to the conclusion, that per foraging bout significantly fewer (nearly halve) flower visits for drinking than for sniffing were executed (Wilcox-Test, p<0.001). It is possible, that many of the flowers did not contain nectar, which is why *E. edwardii* only sniffed at them, but did not drink. Eleven times it was observed, that *E. edwardii* individuals groomed themselves. Either sand baths were carried out, or the rostrum was wiped with the tongue and paws. During this process it was twice observed, that part of the pollen that could be seen on the animals was lost.

*N. famosa* only interacted on two of 16 observation days between 06:50 and 08:58 with *H. hanekomii* inflorescences, which makes them infrequent visitors. When *N. famosa* noticed a *H. hanekomii* inflorescence the animal would fly or hop purposefully towards it, sniff or drink the nectar of the flowers by inserting its beak into the flower opening (Figure 2, C). During this process the stigma was touching the animal’s beak and pollen was deposited on the beaks (Figure 2, D).

A total of 61 flower visits by *N. famosa* were observed where nectar was drunk. Additionally 27 flower visits were observed at which *N. famosa* individuals only sniffed (bringing the beaks in close proximity to the flower entrances). When drinking from a flower the animal stood crouched from the front to the side (90° angle) with its head tilted on the ground in front of the flower. As *N. famosa* drank from the flowers frontally pollen was deposited on the entire beak and on the head up to the eye level. If *N. famosa* drank from the side of the flowers pollen could also be deposited on the beaks and on the sides of the beak. It was observed, that the stigma touched the animals in the same areas. If nectar was present the animal drank from the flowers. If *N. famosa* only briefly stuck its beak into a flower it was probably empty, which the animal might checked by briefly inserting its tongue. In none of the flower visits a destructive behaviour were *N. famosa* injured parts of the corolla or flower organs could be observed. It was also not observed that the animals enlarged the flower entrances. Furthermore no grooming behaviour could be observed. When a flower had nectar and *N. famosa* drank from it the flower visit lasted an average of 1.7s (±1.3s). A wobbling of the inflorescences could never be observed. The frequency with which *N. famosa* took up the nectar could not be determined due to missing visible tongue movement. 12 inflorescence visit by *N. famosa* were observed, with an average time of 14.9s (±13.7s). *N. famosa* only drank once from a flower during an inflorescence visit (Wilcox-Test, p=0.638), and sniffed on a flower only once per inflorescence visit as well (Wilcox-Test, p=0.853). A foraging bout of *N. famosa* was with 39.3s (±24.4s) significant longer than an inflorescence visit of the species lasted on average (Wilcox-Test, p<0.05), which lead to the conclusion that *N. famosa* visited several inflorescences per foraging bout, but came not back to an inflorescence once they changed it (Wilcox-Test, p=0.817). For *N. famosa* 5 foraging bouts from female individuals and 1 sighting without interaction were documented, which lead to an interaction rate of 83.3%. Per foraging bout an average of 12.2 (±7.3) flower visits took place from which nectar was drunk and an average of 5.4 (±4.6) flower visits were made to sniff. Per foraging bout more flower visits could be detected for drinking than for sniffing, which cannot be proven statistically (T-Test, p=0.076). This contradiction is potentially caused by the small dataset, which is present for *N. famosa*. However *N. famosa* only sniffed once per foraging bout on a flower (T-Test, p=0.570) and each flower was only visited once for drinking per foraging bout (T-Test, p=0.570).

Different mice species could be observed between 19:10 and 05:12, as an infrequent flower visitor on *H. hanekomii* inflorescences on 6 observation days. Most of the mice were not interested in the inflorescences and walked near them or searched for food. A few walked purposefully towards the plants and sniffed on the flowers, but never drank from them. When a mouse sniffed on the flowers (Figure 2, E) the animal sometimes used its paws to push the flower entrances apart. It was also found, that the mice nibbled on the corollas, stigmas, styles, and pollen sacs. Only once a mouse was observed cleaning itself by grooming.

It was never observed, that mice were drinking nectar from flowers during an inflorescence visit of *H. hanekomii*. However, 70 flower visits were observed at which mice sniffed. A single inflorescence visit from mice took an average of 10.3s (±7.7s), while the average length of a foraging bout was not significantly longer with 14.9s (±9.2s) (Wilcox-Test, p=0.660), which lead to the conclusion that only a single inflorescence was visited per foraging bout. During a foraging bout a single flower was only visited once for sniffing (Wilcox-Test, p=0.411). Per foraging bout an average of 10.0 (±5.2) flower visits were made to sniff. For mice species 7 foraging bouts and 36 sighting without interaction were documented, which led to an interaction rate of only 16.3 %. It could never be observed, that a stigma touched a mice, even when they were close to the flower openings while sniffing. As the animals often stand head-on to the flowers while they were sniffing the stigmas and pollen sacks were located above the animals, so that pollen could have been deposited on the animals, however a pollen deposition or pollen loads could never be observed on them.

The single inflorescence visits from all observed groups showed no significant differences in length (Figure 3, A). The average length of the foraging bouts between *E. edwardii* [day-night-total] to *N. famosa* as well as from *N. famosa* to Muridae did not differ from each other while E. edwardii [day-night-total] had significantly longer foraging bouts than Muridae (Figure 3, B). *N. famosa* made significantly more flower visits per foraging bout to drink nectar compared to *E. edwardii* [day-night-total], while *E. edwardii* [day-night-total] made significantly more flower visits to drink nectar compared to Muridae (Figure 3, C). The number of flower visits where an animal only sniffed did not differ per foraging bout between all observed groups (Figure 3, D). Flower visits during which nectar was drunk were significantly longer for *E. edwardii* compared to *N. famosa* (Figure 3, E). Additionally flower visits during which nectar was drunk were significantly longer for *E. edwardii* in the dark compared to *E. edwardii* at light (Figure 3, E).

**Figure 3:**
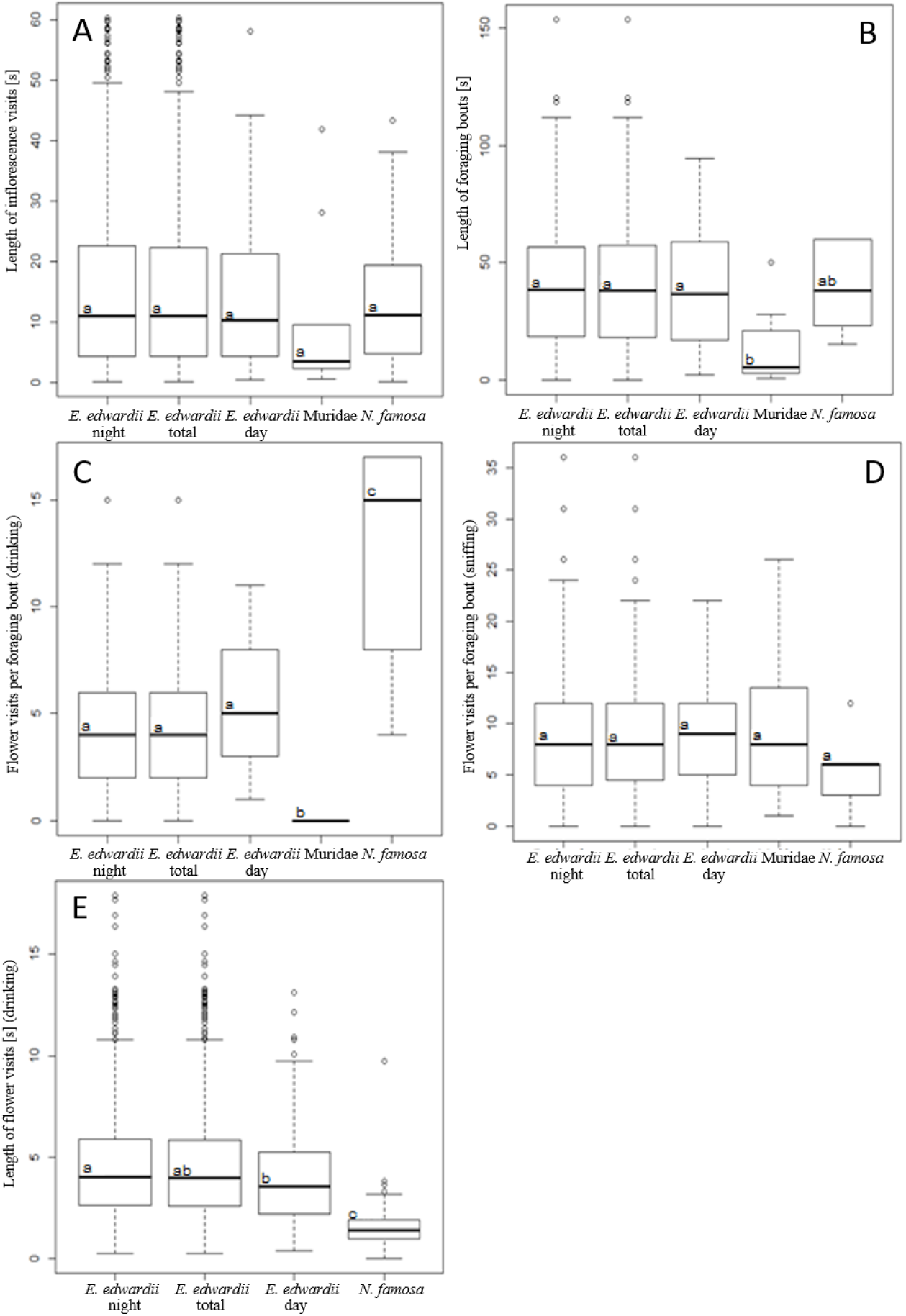
Interaction of different species with *H. hanekomii*. **A:** Length of inflorescence visits. Kruskal-Wallis: Chi^2^=2.906; f=4; p=0.574. **B:** Length of foraging bouts. Kruskal-Wallis: Chi^2^=7.705; f=4; p=0.103. **C:** Number of flower visits per foraging bout were nectar was drunk. Kruskal-Wallis: Chi^2^=28.304; f=4; p=0.001. **D:** Number of flower visits per foraging bout where animals only sniffed. Kruskal-Wallis: Chi^2^=1.897; f=4; p=0.755. **E:** Length of flower visits during which nectar was drunk. Kruskal-Wallis: Chi^2^=102.380; f=4; p=0.001. For each panel the middle line represents the median, the upper and lower line represent the quartiles, and the satellites cover the whole measuring range. Outlier are not included and are displayed as separate dots. Boxes with the same letters do not differ significantly. The data is based on attachment Table 1 – 6.

As only *E. edwardii* and *N. famosa* seem to feed on *H. hanekomii* the focus for pollen deposition and stigma touches were placed on those species. During flower visits when *E. edwardii* and *N. famosa* drank nectar, it was often observed that the stigma touched the animals and that pollen was deposited on them (Figure 4). For *E. edwardii* most stigma touches occurred above or on the rostrum, but stigma touches also occurred between the eyes, on the forehead, on the cheeks, on the trunk and on its sides. If there was no contact between the stigma and the animal *E. edwardii* would often crouch from below, or from the sides with its head tilted, so that the stigma was next to the animal or above it causing a great distance which prohibited contact. Most pollen deposition areas coincided with the areas of stigma touches. If no pollen deposition could be observed *E. edwardii* often stood in such a way that no statement could be made as to whether pollen was deposited or not. Alternatively in those cases insufficient pollen amounts were deposited on *E. edwardii* since only large amounts of pollen were recognizable in the videos, or the anthers were already empty. In case of *E. edwardii* it was often observed that after visiting flowers more pollen was seen on the animals than before even if no active deposition was observed. Despite matching locations of pollen deposition and stigma contact on *E. edwardii* it could only be observed once that the stigma actually touched deposited pollen by pressing it on the fur on the rostrum on which pollen was located. This discrepancy could be caused by the large number of interactions were either stigma touches or pollen deposition or none of both could be observed due to the position of the animal between inflorescence and camera. For *N. famosa* the places where pollen was deposited and where the stigma touched the animals coincided as well. The most stigma touches were observed on the beak and the head of the animal, but could also be detected on the side of the beak and at the beak attachment point. Pollen deposition was observed in the same areas. If no stigma touches or pollen deposition occurred *N. famosa* stood heads on or from the sides to the inflorescences causing stigma and pollen sacs to be a few millimetres to the side of the beak, so that no contact or deposition was possible. Despite matching locations of pollen deposition and stigma contact on *N. famosa* it could never be observed that the stigma actually touched deposited pollen, which was potentially caused by the small data set present for this species. In case of *N. famosa* it was observed as well that after visiting flowers more pollen could be seen on the animals than before even if no active deposition was observed.

**Figure 4:**
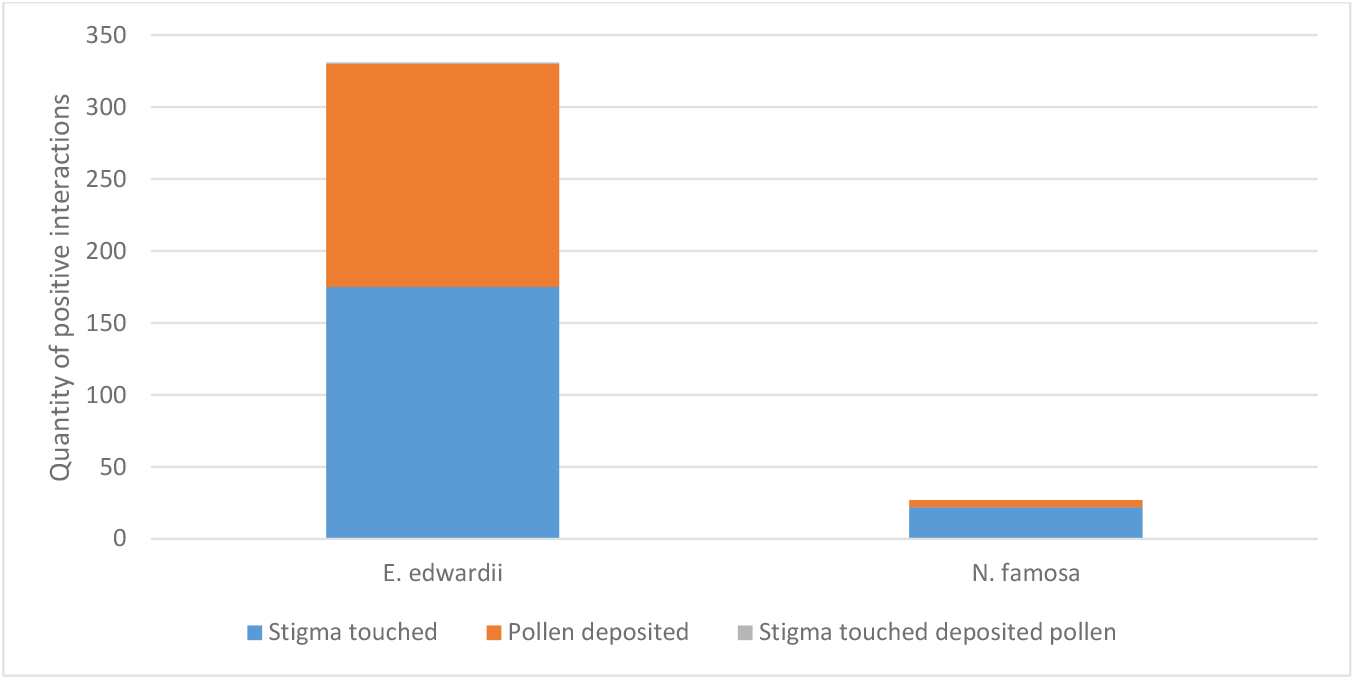
Frequency of stigma touches and pollen deposition of *H. hanekomii* inflorescences on *E. edwardii* and *N. famosa* after they drank from the flowers. For *E. edwardii* 1031 and for *N. famosa* 61 flower visits were recorded representatively. The differences result from the number of flower visits where it was not visible whether pollen was deposited or the stigma touched the animal. The data is based on attachment Table 1.

If only *E. edwardii* and *N. famosa* are considered as pollinators, due to non-destructive flower interaction, potential pollen transfer, nectar drinking and sniffing on flowers and if drinking and sniffing are considered as flower interactions 97.3 % of all interactions in the whole observation period are executed by *E. edwardii* and only 2.7 % by *N. famosa* (Figure 5, left). This numbers change slightly to 80 % interaction of *E. edwardii* and 20 % interaction of *N. famosa*, if only the day period is observed (Figure 5, right).

**Figure 5:**
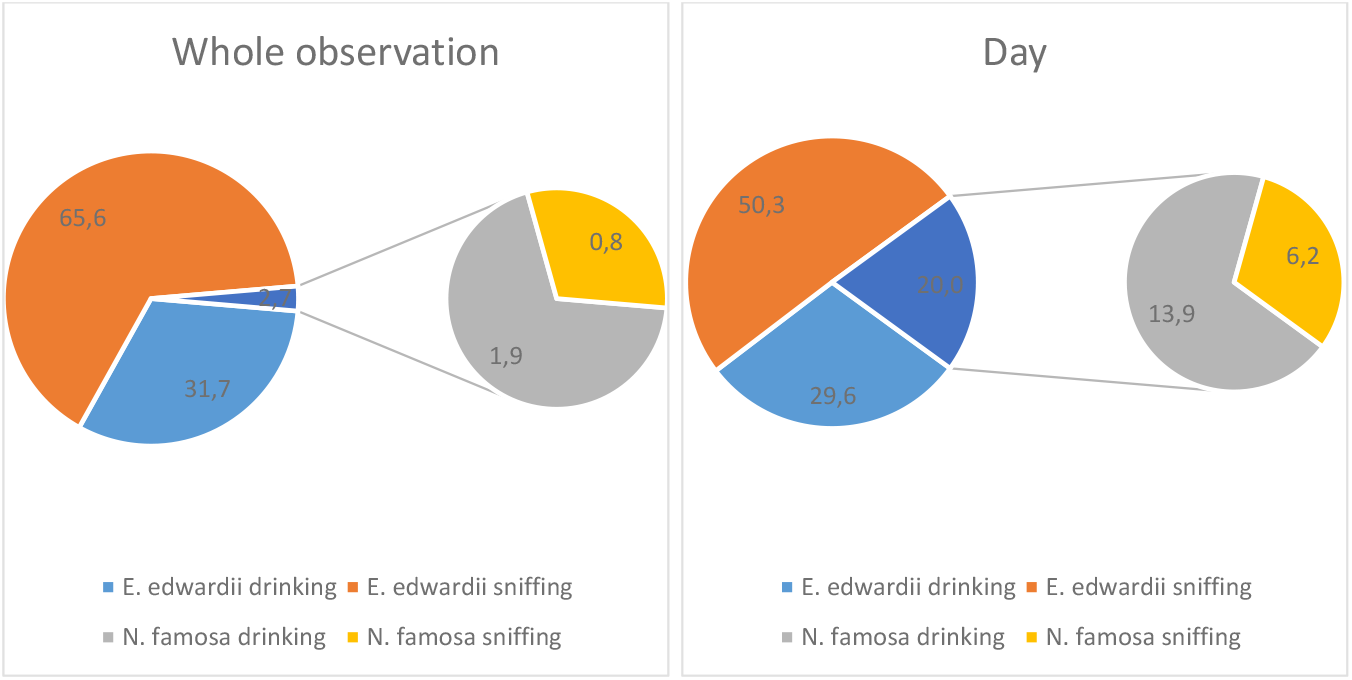
Relative ratio of flower interactions on *H. hanekomii* inflorescences by *E. edwardii* and *N. famosa* **Left:** Whole observation periode. **Right:** Only observations in daylight. The data is based on attachment Table 1.

UV photography of *H. hanekomii* inflorescences (Figure 6, A-A”) and single flowers (B-B”) showed no UV reflecting spots. No stinging hairs could be observed on the flowers (Figure 6, C-C”). The lack of UV reflecting spots on single flowers was proven by spectrometry as well (Figure 7). The flowers show no reflection in the UV spectrum. They only reflected light between 600nm and 850nm.

**Figure 6:**
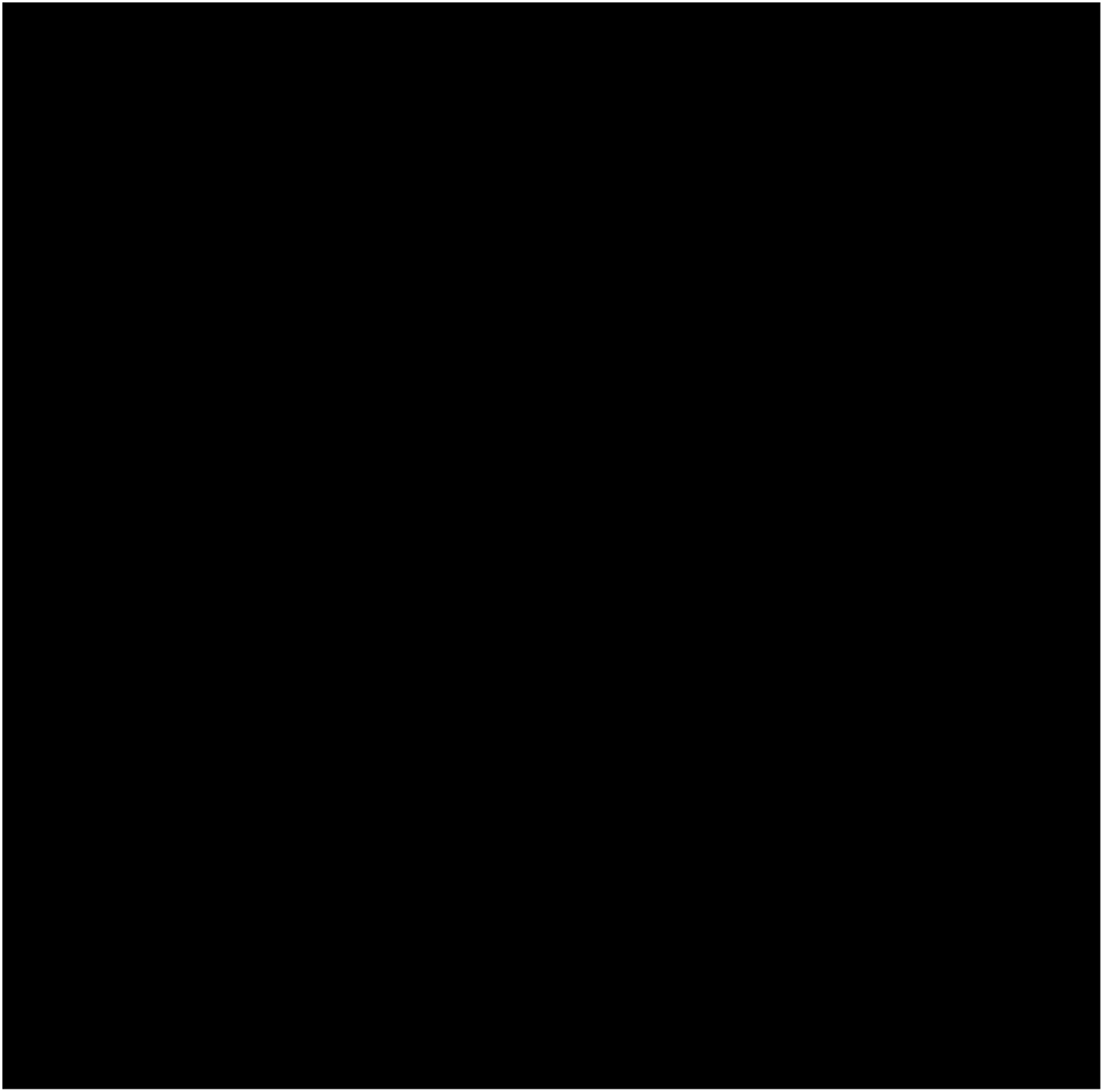
Images of *H. hanekomii* Inflorescences and flowers. **A:** *H. hanekomii* inflorescence in the field. A’ resembles the same plant photographed without an UV filter to show UV reflecting areas, while A” is additionally illuminated with an UV lamp. **B:** *H. hanekomii* flowers. B ‘ ‘ resembles the same flowers photographed without an UV filter to show UV reflecting areas, while B” is additionally illuminated with an UV lamp. **C:** Hair on *H. hanekomii* flowers in 50× magnification. C’ resembles the same flower in 200× magnification and C” resembles the same flower in an unknown magnification. Pictures taken by Niedzwetzki-Taubert, T., Karvang, A.S.N., Wester, P. **Pictures not shown due to copyright reasons**

**Figure 7:**
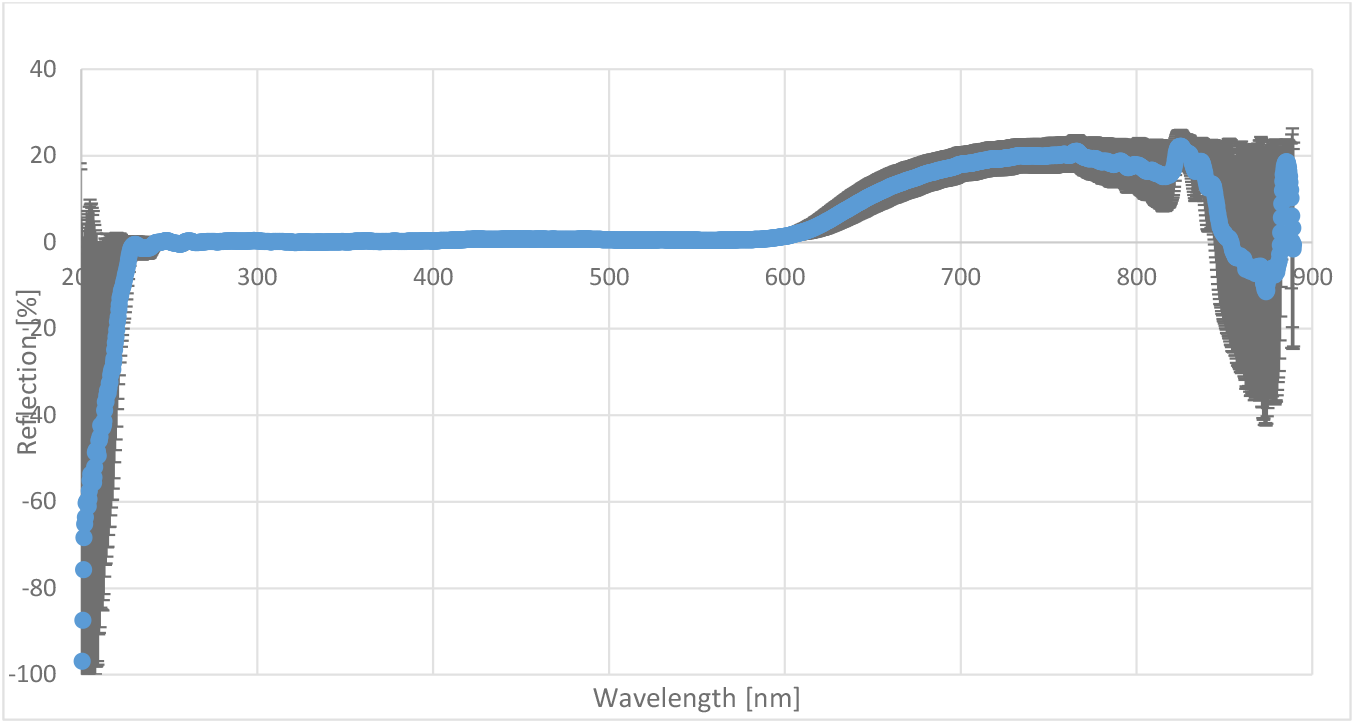
JAZ spectum of H. hanekomii flowers. The average and standard deriviation of 32 flowers are shown in the UV to infrared wavelength. As a measuring device the JAZ USB Spectrophotometer (Ocean Optics, Dunedin, USA) was used.

The flower entrance width and height are significantly smaller for *H. hanekomii* than for *H. atropurpurea* (Figure 8). Other flower characteristics show significant differences as well, but can only be evaluated in combination with the beak and rostrum sizes of the animals. Plant height, amount of flowers, and corolla length show no significant differences between the two species.

**Figure 8:**
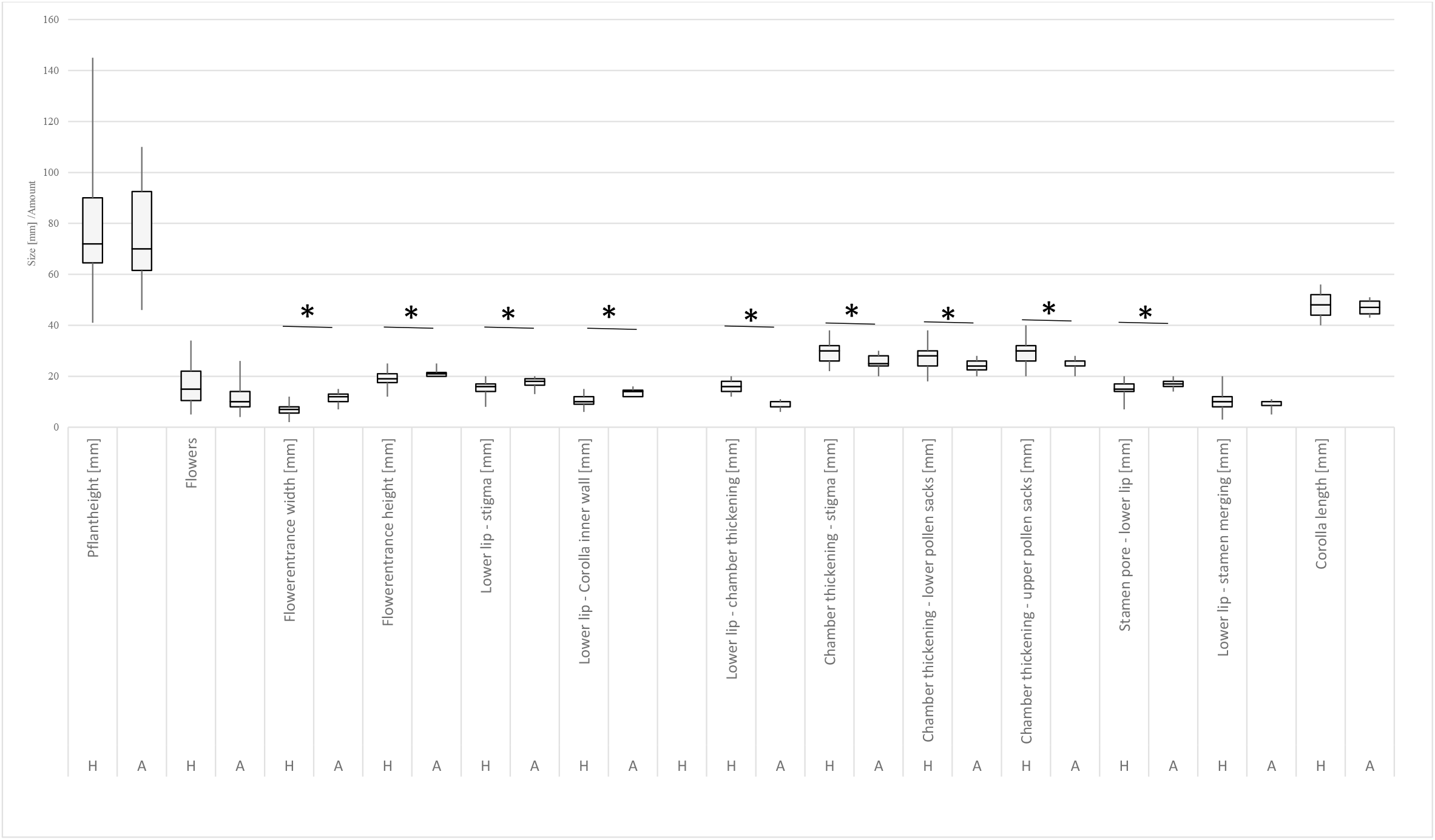
Morphological comparison between *H. hanekomii* (H, n=31) and *H. atropurpurea* (A, n=11) for different plant traits. For each panel the middle line represents the median, the upper and lower line represent the quartiles, and the satellites cover the whole measuring range. Boxes connected with an asterisk show a significant difference (T-Test, p<0.05, two sided, heteroscedastic)

## Discussion

*H. hanekomii* is pollinated by *E. edwardii* and *N. famosa*. However, the pollination by *N. famosa r*equires further research.

Chersina angulata, Papio ursinus, and Bos primigenius taurus can be excluded as pollinators, as they only display destructive behaviour on *H. hanekomii*. Insects can also be ruled out as pollinators, since they were not seen on *H. hanekomii* inflorescences during the entire observation period. Muridae can be excluded as pollinators of *H. hanekomii* as well, since they only sniffed on flowers when visiting them, did not drank the nectar (Figure 2, E), no pollen was deposited, or the stigma touched the animals, and they showed destructive behaviour on the flowers, what a pollinator should not exhibit. Muridae are known as pollinators of other plants, on which they drink the sugar rich nectar and use it as a food source (Biccard and Midgley, 2009; Johnson *et al.*, 2001, 2011; Letten and Midgley, 2009; Rourke and Wiens, 1978; Turner *et al.*, 2011; Wester, 2015; Wester *et al.*, 2009). As they do not consume the nectar of *H. hanekomii* it is unlikely that they act as a pollinator of the species. Potentially the flower entrances were to small and tubular for their heads so that they could not enter them, which was already noted for *H. atropurpurea* by Wester (2011).

The data proves that *E. edwardii* in its natural environment voluntarily and regularly targeted *H. hanekomii* inflorescences, to forage nectar on their flowers. During this process usually several flowers and inflorescences were visited one after another. The targeted foraging on *H. hanekomii* and the fact that several inflorescences are visited every night leads to the conclusion that *E. edwardii* is a pollinator of *H. hanekomii*. A pollinator must visit the flowers regularly, which is achieved by a direct attractant (Faegri & Van der Pijl, 1979), in this case nectar. The fact that > 95 % of all flower visits were made by *E. edwardii* supports this conclusion (Figure 5). A plant is considered to be adapted to a functional group of pollinators if > 75 % off all interactions are with that group (Fenster *et al.*, 2004), which applies for *E. edwardii*. Thus *H. hanekomii* is adapted to pollination by *E. edwardii* and this species is most likely the main pollinator. Many of the points that indicate whether a species is pollinated by non-flying mammals apply here (Johnson *et al.*, 2001; Rourke and Wiens, 1977, 1978; Wiens *et al.*, 1983). No flowers or other parts of the plant were damaged during any of the flower visits by *E. edwardii*. Pollen could be detected on the animals, which was deposited on the snouts when drinking from the flowers, as well as the pollen transport from flower to flower. Contact between the deposited pollen and the stigma could also be observed. Furthermore an accurate fit between the snouts of the animals and the flowers could be determined. Missing stinging hairs and UV reflection spots, as well as the colours spectrum of the plant support this hypothesis as well.

Although *N. famosa* rarely visited *H. hanekomii* inflorescences this species can be considered as a pollinator as well. Only five *N. famosa* individuals were observed drinking nectar from *H. hanekomii*. Despite this small number of sightings it could be observed that the individuals voluntarily and purposefully went to the plants to drink nectar without damaging them (Figure 2, C). Pollen deposition and stigma contacts could also be observed for *N. famosa*, which speaks for this species as a pollinator. No contact between the stigma and on the animal deposited pollen could be observed, which is most likely caused by the low quantity of sightings. The small number of sightings of *N. famosa* might be caused in natural fluctuations in the population density, but is very low compared to *E. edwardii*. Only 2.7 % of all flower visits are made by *N. famosa* which makes this species a secondary pollinator of *H. hanekomii* next to *E. edwardii* as the main pollinator.

Several artefacts need to be considered which may have resulted in the low amount of *N. famosa* sightings.

The first reason is a Protea farm (Rockwood) located less than a kilometre southeast of the *H. hanekomii* population. Bird pollinated Proteaceae grow on this farm and were visited by sunbirds including *N. famosa* (personal observation). As the *H. hanekomii* inflorescences grew at ground level, cryptic under bushes they are less showy than the outnumbering Proteaceae on the farm which have large exposed showy floriferous inflorescences. Suggesting that the sunbirds were more focused on the Proteaceae as a more lucrative food source and therefore did not visit *H. hanekomii* inflorescences to drink nectar because they were far less attractive at ground level. Another reason for the low number of *N. famosa* observations could be that the birds were too scared to forage on the plants. Birds operate at a flight distance, which is the distance at which a bird will flee from a thread (Battle *et al.*, 2016). Additionally the alarm distance describes the distance between a bird and a source of disturbance from which the animal changes its natural behaviour (Ruddock and Whitfield, 2007). It is possible, that the cameras were positioned too close to the plants and were located within the flight or alert distance of the birds and therefore only those individuals who had smaller escape or alert distances interacted with the plants and did not feel disturbed by the cameras. Possibly more birds would seek out *H. hanekomii* inflorescences if the cameras were placed further away (which is possible with long distance lenses) or camouflaged. In addition *N. famosa* unlike *E. edwardii* is diurnal (Wellmann and Downs, 2009). However the cameras are more noticeable in daylight and could be more disruptive which may have deterred animals. Camera traps have not been widely used to identify pollinators and if yes it was mostly done for rodent pollinated species but could also be applied to bird pollinated species (Melidonis and Peter, 2015; Zoeller *et al.*, 2016), which was proven by (Turner and Midgley, 2016) and represents a more efficient and discrete way of documenting flower visits by larger animals than field observations by humans, which might disturb the animals (Zoeller *et al.*, 2016). Thereby camera traps open up the possibility of documenting natural interactions of unstressed animals as they do not disturb the foraging species whether rodents or birds (Melidonis and Peter, 2015; Zoeller *et al.*, 2016). However the possibility that the animals were disturbed by the cameras cannot be ruled out, as it is possible that in these works the cameras were located further away or where better camouflaged or the video material was not gathered as frequently and therefore did not affect the animals. The last reason why *N. famosa* may have avoided the *H. hanekomii* inflorescences is a change in the environment. Sunbirds often perch on various vegetative or reproductive plant structures during foraging as they prefer exposed positions to get a better view of their surrounding (Turner and Midgley, 2016; de Waal *et al.*, 2012). Not all sunbird pollinated plants possess such structures. Some examples are *Babiana carminea, Cytinus sanguineus, Lachenalia luteola*, and *H. sanguinea* (Hobbhahn and Johnson, 2015; Turner and Midgley, 2016; de Waal *et al.*, 2012). Sunbirds foraging on these plants must sit on the ground to drink nectar. According to Hobbhahn & Johnson (2015) there is a possibility that birds visit geofloral flowers without perches, as the surrounding bushes provide a sense of security and the birds do not feel threatened by predators when foraging. *H. hanekomii* grows under bushes similar to *H. sanguinea* and does not possess perches. In order to record better with the camera traps the environment around the *H. hanekomii* inflorescences was modified by removing shrubs and other plants around them (~working distance of the camera traps). This procedure might be the reason why the birds did not feel safe around the plants and therefore did not visit them. This is also supported by the fact that most *N. famosa* individuals foraged on *H. hanekomii* inflorescences in the vicinity of which there were still protective bushes or stones.

The fact that the length of inflorescence visits between *E. edwardii* [day-night-total], *N. famosa* and Muridae did not differ from each other shows that each species interacts with the inflorescences for a similar length of time regardless their intention and independently of the time of the day (Figure 3). The fact that the foraging bouts of *E. edwardii* and *N. famosa* are significantly longer than those of Muridae (Figure 3) shows that those species visit more inflorescences per foraging bout than Muridae which further increases their importance as pollinators, since they distribute the pollen between different plants of one population. The fact that no difference could be detected between *E. edwardii* in the light and in the dark or between *E. edwardii* in the light and *N. famosa* with regard to the length of the foraging bout proves that the brightness also has no influence on how long the animals look for food on the plants (Figure 3). Since *N. famosa* visits highly significantly more flowers to drink nectar from than *E. edwardii* during a foraging bout this may compensate the less frequent visits (Figure 3). *E. edwardii* drinks from flowers significantly longer than *N. famosa* (Figure 3). *E. edwardii* might feel more secure on the plants at night and therefore drink longer from the flowers, which is supported by the fact, that *E. edwardii* flower visits for drinking are significantly shorter at day than at night (Figure 3). However *E. edwardii* flower visits are still significantly longer at day than the visits of *N. famosa*, which indicates that either this species feels safer on the plants or, that *N. famosa* has a better access to the nectar allowing him to drink more or faster. This cannot be proven, as no frequency of licking could be determined for *N. famosa*. Alternatively *N. famosa* may drink from the flowers shorter so that the animal can better observe its surroundings during the day, as the observed sunbirds generally seemed more nervous than the observed elephant shrews.

There are several possible explanations why *H. hanekomii* is pollinated by birds and non-flying mammals.

The first is, that *H. hanekomii* is a species in transmission from avian to mammalian pollination or vice versa. Rourke & Wiens (1977) have already hypothesised that some Proteaceae which are pollinated by non-flying mammals have evolved from bird pollinated species. According to Wiens et al. (1983) a switch from one pollination syndrome to another occurs when one pollinator group has several advantages over the other pollinator group. For example, if they are more reliable pollinators of a species by transferring more pollen or by being more concentrated in the area. Another example of a change in a pollinator system was observed as bees can replace sunbirds as effective pollinators (Paton, 2000) and vice versa (Strelin *et al.*, 2016). All these examples show that a pollination system is not rigid, and that the pollinator of a species can change due to different factors such as populations or competitors. Assuming that *H. hanekomii* is an originally bird pollinated species (due to its morphological similarity to *H. sanguinea*, the striking red flowers, the lack of fragrance, and the small flower openings) it can be assumed that a switch to mammalian pollination is taking place as sunbirds are no longer reliable pollen carriers. Reasons for this could be stagnant sunbird populations or competition between *H. hanekomii* and the Proteaceae on the nearby farm. As only *N. famosa* was sighted on the plants drinking nectar it can be concluded that *H. hanekomii* is specialised for long beaked sunbird species and belongs to the malachite sunbird pollination syndrome which was described by Geerts & Pauw (2009). This is supported by the fact that no nectar stealing was observed which was already observed for other sunbird species on other plants (Vogel, 1954). A pollinator switch from mammal to avian pollination is rather unlikely given the abundance of *E. edwardii* individuals interacting with the plants. If *H. hanekomii* is currently in a transitional stage between ornithophilic and non-flying mammal pollination syndromes this could explain the extraordinary high variance within the species (Figure 1, A & B). Some individuals may have already evolved a wider flower entrance to allow better entry for mammals, while other individuals may not have completed this step (Figure 1, D & E). Another possibility why *E. edwardii* and *N. famosa* were sighted on the inflorescences could be that *H. hanekomii* is not specialised for one functional group as pollinators and is pollinated by several groups. In this case a mixed pollination system would be present (Queiroz *et al.*, 2016; Ramírez-Aguirre *et al.*, 2016), where *N. famosa* and *E. edwardii* contribute to pollination if traits from both pollination syndromes are present, which is the case. In that case *H. hanekomii* would had adapted to several functional groups which would also lead to the morphological variance that was observed (Figure 1, A-E). Adaption to *N. famosa* and *E. edwardii* would result in long corollas (Figure 1, D & E), with distances from the flower entrance to the nectar being about the animals rostrum or beak length, excluding short snouted mice or birds as pollinators.

Depending on which pollinator group *H. hanekomii* adapts evolutionary processes and plant traits are driven in one or the other direction. For example the wide colour spectrum (Figure 1, A & B) could increase if *H. hanekomii* further adapts to *E. edwardii* as a pollinator since they see dichromatic (Thüs *et al.*, 2020) and cannot differentiate the red colour of the plants. The scent on the other hand could shift from being odourless to a nutty, buttery smell like found in other *E. edwardii* pollinated Hyobanche species (Wester, 2011) as the scent plays an important role to attract those animals to flowers. If *H. hanekomii* would further adapt to *N. famosa* as a pollinator the colour spectrum would likely decrease as more intense red colorations would be an advantage to attract birds. Additionally the scent of the plants should not change since it does not play a role in bird pollinated plant species. If *H. hanekomii* adapts to both pollinator groups the plant traits would reach an intermediate state between the two pollination syndromes, so that the plants are adapted to both groups.

UV photography of *H. hanekomii* inflorescences and single flowers (Figure 6, A-A” and B-B”) support the hypothesis of adaption to non-flying mammal pollination as no UV reflecting spots could be detected that would attract birds or insects. This observation is supported by a lack of stinging hairs as well (Figure 6, C-C”). The lack of UV reflecting spots on single flowers could be proven by spectrometry as well (Figure 7). The flowers show no reflection in the UV spectrum at all. They only reflect light between 600nm to 850nm which correlates perfectly with their red to dark red colour. In contrary to this some of the most important flower morphological characteristics indicate bird pollination. The flower entrance width and height are significantly smaller for *H. hanekomii* than for *H. atropurpurea* (Figure 8) which is indicating a better adaption to birds with narrow beaks or elephant shrews excluding all rodents from the plants. This would explain their destructive behaviour as well. However definite statements could only be given here, if the plants are compared to *H. sanguinea* and to the beaks and rostra of the potential pollinators.

## Outlook

Video observations should be repeated on a different population which is not close to a huge amount of Proteaceae to test if more *N. famosa* individuals visit *H. hanekomii* under these conditions since they could play a bigger role in pollination than indicated in this work. Additional the cameras should be camouflaged better, and not be visited that often, and the surrounding of the plants should not be artificially altered.

However no conclusive statement can be made about the unique pollinator since the pollinator efficiency of *E. edwardii* and *N. famosa* was not examined. This could be analysed by excluding certain pollinator groups from the plants and analysing the Seed set as it was executed for *H. sanguinea* by Turner & Midgley (2016). This could prove which species is the more efficient pollinator.

Furthermore *H. hanekomii* nectar should be analysed to determine its sugar content and composition. As there is a connection between the sugar composition of nectar and the pollinator (Baker and Baker, 1983). This data could be compared to data from known bird and mammal pollinated plants. Sunbird pollinated plants contain sucrose rich nectar (40 % - 60 %) (Johnson and Nicolson, 2008), while nectar from non-flying mammal pollinated plants vary in their sugar composition (Wester *et al.*, 2016). In combination to this the amount of nectar should be monitored during the day to determine whether more nectar is produced during the day or at night, like it was suggested by (Wiens *et al.*, 1983).

In addition snout, head and beaks swabs from the animals should be collected to compare the amount of pollen the different species transport between plants. Samples of faeces should be examined as well in order to determine how much pollen gets into the animals stomachs through cleaning or eating.

Also the scent of *H. hanekomii* should be analysed, for example by GC-MS like it was done by Wester et al. (2016) since it is known that scents are important attractants to lure pollinators to flowers (Williams, 1983). As a lack of odour is rather atypical for plants pollinated by non-flying mammals, but typical for bird pollinated plants (Wester *et al.*, 2016). Chemical substances such as aliphatic ketones in the nectar can be attractive to non-flying mammals, while in contrast benzenoid compounds are present in the nectar of bird pollinated plants (Johnson *et al.*, 2011; Steenhuisen *et al.*, 2012). The chemical substances present in *H. hanekomii* nectar could therefore give a statement if the scent is more attractive to sunbirds or elephant shrews.

## Acknowledgements

I hereby thank Prof. Dr. Lunau for the possibility to conduct this research in the Institute for sensory ecology at the Heinrich-Heine-University Düsseldorf and Dr. Wester for the help with fieldwork as well as for the supervision of the bachelor thesis which it the foundation of this work and the organisation of the necessary permits. [INSERT PERMITS HERE]. I thank the owners of the Fonteintjie Farm that we were able to conduct research on their property. I thank the Wolters-Vollhardt-foundation for the scholarship of Tim Niedzwetzki-Taubert which enabled the research stay in South Africa. I thank Sebastian Köthe for the kind help in terms of statistical analysis. Finally I thank Anna Sophia Natascha Karvang for all her support, help with fieldwork, help with discussions about topics of this publication and so much more.

