## Supplement 1 for "*Hyobanche hanekomii* (Orobanchaceae) is pollinated by non-flying mammals and birds"

**Table 1:** Behavior of observed animal species during video observation of *H. hanekomii* inflorescences. Number of flowers visited per foraging bout and inflorescence visit (drinking or sniffing). Length of the respective inflorescence visits, foraging bouts and drinking actions. Pollen Deposition on the animals and damage to the flowers.

| | | *E. edwardii* | *N. famosa* | Muridae |
|---|---|---|---|---|
| **Days on which the animal is observed** | | 16 | 3 | 15 |
| **Days with foraging bouts** | | 16 | 2 | 6 |
| **Time of activity** | | 18:16-07:58 | 06:50-08:58 | 19:10-05:12 |
| **Sum of visited inflorescences** | Total | 382 | 8 | 10 |
| | Day | 43 | 8 | 0 |
| | Night | 339 | 0 | 10 |
| **Average number of visited inflorescences per foraging bout and their minimum / maximum number** | Total | 1.566 (±0.904) | 1.600 (±0.816) | 1.429 (±0.229) |
| | | 1 to 4 | 1 to 2 | 1 to 2 |
| | Day | 1.720 (±0.793) | 1.600 (±0.816) | 0.000 (±0.000) |
| | | 1 to 3 | 1 to 2 | 0 |
| | Night | 1.548 (±0.800) | 0.000 (±0.000) | 1.429 (±0.229) |
| | | 1 to 4 | 0 | 1 to 2 |
| **Sum of inflorescence visits** | Total | 548 | 12 | 10 |
| | Day | 64 | 12 | 0 |
| | Night | 484 | 0 | 10 |
| **Average number of inflorescence visits per foraging bout and their minimum / maximum number** | Total | 2.246 (±1.670) | 2.400 (±2.098) | 1.429 (±0.229) |
| | | 1 to 8 | 1 to 6 | 1 to 2 |
| | Day | 2.560 (±1.718) | 2.400 (±2.098) | 0.000 (±0.000) |
| | | 1 to 7 | 1 to 6 | 0 |
| | Night | 2.210 (±1.616) | 0.000 (±0.000) | 1.429 (±0.229) |
| | | 1 to 8 | 0 | 1 to 2 |
| **Average length of inflorescence visits [s] and their minimum / maximum length [s]** | Total | 15.582 (±14.502) | 14.883 (±13.672) | 10.278 (±7.717) |
| | | 0.183 to 60.268 | 0.217 to 43.347 | 0.600 to 41.900 |
| | Day | 15.509 (±11.849) | 14.883 (±13.672) | 0.000 (±0.000) |
| | | 0.533 to 44.167 | 0.217 to 43.347 | 0 |
| | Night | 19.628 (±17.028) | 0.000 (±0.000) | 10.278 (±7.717) |
| | | 0.183 to 60.268 | 0 | 0.600 to 41.900 |
| **Sum of flower visits (nectar drinking)** | Total | 1031 | 61 | 0 |
| | Day | 130 | 61 | 0 |
| | Night | 901 | 0 | 0 |
| **Mean number of flower visits per inflorescence (nectar drinking) and their minimum / maximum number** | Total | 1.881 (±1.821) | 5.083 (±4.733) | 0.000 (±0.000) |
| | | 0 to 9 | 1 to 17 | 0 |
| | Day | 2.464 (±1.835) | 5.083 (±4.733) | 0.000 (±0.000) |
| | | 0 to 7 | 1 to 17 | 0 |
| | Night | 2.435 (±2.097) | 0.000 (±0.000) | 0.000 (±0.000) |
| | | 0 to 9 | 0 | 0 |
| **Sum of flowers visited (nectar drinking)** | Total | 989 | 51 | 0 |

| | | | | |
|---|---|---|---|---|
| | Day | 130 | 51 | 0 |
| | Night | 859 | 0 | 0 |
| **Mean number of flowers visited per inflorescence (nectar drinking) and their minimum / maximum number** | Total | 1.805 (±1.708) | 4.250 (±3.883) | 0.000 (±0.000) |
| | | 0 to 9 | 1 to 14 | 0 |
| | Day | 2.464 (±1.835) | 4.250 (±3.883) | 0.000 (±0.000) |
| | | 0 to 7 | 1 to 14 | 0 |
| | Night | 2.297 (±1.932) | 0.000 (±0.000) | 0.000 (±0.000) |
| | | 0 to 9 | 0 | 0 |
| **Sum of flower visits (sniffing)** | Total | 2130 | 27 | 70 |
| | Day | 221 | 27 | 0 |
| | Night | 1909 | 0 | 70 |
| **Mean number of flower visits per inflorescence (sniffing) and their minimum / maximum number** | Total | 3.887 (±3.360) | 2.250 (±3.523) | 7.000 (±0.688) |
| | | 0 to 21 | 0 to 12 | 1 to 20 |
| | Day | 3.929 (±3.431) | 2.250 (±3.523) | 0.000 (±0.000) |
| | | 0 to 13 | 0 to 12 | 0 |
| | Night | 4.494 (±3.918) | 0.000 (±0.000) | 7.000 (±0.688) |
| | | 0 to 21 | 0 | 1 to 20 |
| **Sum of flowers visited (sniffing)** | Total | 1584 | 18 | 47 |
| | Day | 170 | 18 | 0 |
| | Night | 1414 | 0 | 47 |
| **Mean number of flowers visited per inflorescence (sniffing) and their minimum / maximum number** | Total | 2.891 (±2.155) | 1.500 (±2.063) | 4.700 (±0.688) |
| | | 0 to 10 | 0 to 6 | 1 to 10 |
| | Day | 2.964 (±2.317) | 1.500 (±2.063) | 0.000 (±0.000) |
| | | 0 to 9 | 0 to 6 | 0 |
| | Night | 3.255 (±2.425) | 0.000 (±0.000) | 4.700 (±0.688) |
| | | 0 to 10 | 0 | 1 to 10 |
| **Average length of the foraging bouts [s] and their minimum / maximum length [s]** | Total | 38.235 (±27.102) | 39.295 (±24.407) | 14.883 (±9.175) |
| | | 0.148 to 153.567 | 15.283 to 60.050 | 0.600 to 50.234 |
| | Day | 38.533 (±25.784) | 39.295 (±24.407) | 0.000 (±0.000) |
| | | 2.150 to 94.551 | 15.283 to 60.050 | 0 |
| | Night | 38.201 (±26.118) | 0.000 (±0.000) | 14.883 (±9.175) |
| | | 0.148 to 153.567 | 0 | 0.600 to 50.234 |
| **Number of foraging bouts“** | Total | 244 | 5 | 7 |
| | Day | 24 | 5 | 0 |
| | Night | 220 | 0 | 7 |
| **Sightings without foraging bouts** | Total | 53 | 1 | 36 |
| | Day | 4 | 1 | 0 |
| | Night | 49 | 0 | 36 |
| **Mean number of flower visits per foraging bout (nectar drinking) and their minimum / maximum number** | Total | 4.225 (±3.030) | 12.200 (±7.250) | 0.000 (±0.000) |
| | | 0 to 15 | 4 to 17 | 0 |
| | Day | 5.200 (±3.096) | 12.200 (±7.250) | 0.000 (±0.000) |

| | | | | |
|---|---|---|---|---|
| | | 1 to 11 | 4 to 17 | 0 |
| | Night | 4.114 (±2.776) | 0.000 (±0.000) | 0.000 (±0.000) |
| | | 0 to 15 | 0 | 0 |
| **Mean number of visited flowers per foraging bout (nectar drinking) and their minimum / maximum number** | Total | 4.053 (±2.894) | 10.200 (±5.925) | 0.000 (±0.000) |
| | | 0 to 15 | 3 to 14 | 0 |
| | Day | 5.200 (±3.200) | 10.200 (±5.925) | 0.000 (±0.000) |
| | | 1 to 11 | 3 to 14 | 0 |
| | Night | 3.922 (±2.656) | 0.000 (±0.000) | 0.000 (±0.000) |
| | | 0 to 15 | 0 | 0 |
| **Mean number of flower visits per foraging bout (sniffing) and their minimum / maximum number** | Total | 8.730 (±6.324) | 5.400 (±4.550) | 10.000 (±5.168) |
| | | 0 to 36 | 0 to 12 | 1 to 26 |
| | Day | 8.840 (±4.871) | 5.400 (±4.550) | 0.000 (±0.000) |
| | | 0 to 22 | 0 to 12 | 0 |
| | Night | 8.717 (±6.038) | 0.000 (±0.000) | 10.000 (±5.168) |
| | | 0 to 36 | 0 | 1 to 26 |
| **Mean number of visited flowers per foraging bout (sniffing) and their minimum / maximum number** | Total | 6.492 (±4.548) | 3.600 (±2.683) | 6.714 (±3.241) |
| | | 0 to 28 | 0 to 6 | 1 to 16 |
| | Day | 6.800 (±3.478) | 3.600 (±2.683) | 0.000 (±0.000) |
| | | 0 to 13 | 0 to 6 | 0 |
| | Night | 6.457 (±4.352) | 0.000 (±0.000) | 6.714 (±3.241) |
| | | 0 to 28 | 0 | 1 to 16 |
| **Mean length of flower visits (nectar drinking) [s] and their minimum / maximum length [s]** | Total | 4.495 (±2.860) | 1.681 (±1.330) | 0.000 (±0.000) |
| | | 0.283 to 17.867 | 0.034 to 9.733 | 0 |
| | Day | 3.735 (±2.429) | 1.681 (±1.330) | 0.000 (±0.000) |
| | | 0.967 to 8.733 | 0.034 to 9.733 | 0 |
| | Night | 4.459 (±3.004) | 0.000 (±0.000) | 0.000 (±0.000) |
| | | 0.283 to 17.867 | 0 | 0 |
| **Average nectar licking frequency [Hz] and their minimum / maximum number** | | 4.332 (±1.766) | Not detectable | No drinking |
| | | 2 to 6 | Not detectable | No drinking |
| **Does the animal reach the nectar?** | | Yes, if a flower contains nectar the animal drinks. If the flower is empty this is tested by inserting the tongue briefly or by sniffing. | Yes, if a flower contains nectar the animal drinks. If the flower is empty this is tested by inserting the beak briefly. | No |
| **Does the stigma touch the animal and if so where?** | | Not detectable: 829 | Not detectable: 29 | No |
| | | Over or on the snout: 136 | head: 6 | |
| | | Between the eyes: 2 | On the beak: 8 | |
| | | On the forehead: 12 | Sides of the beak: 3 | |
| | | Nose: 8 | Beak attachment point: 2 | |
| | | Side of the nose:14 | Sides of the beak & beak: 3 | |
| | | Cheek:4 | No: 10 | |
| | | No: 26 | | |
| **Is pollen deposited on the animal and if so where?** | | Not detectable: 817 | Not detectable: 54 | No |

| | | | |
|---|---|---|---|
| | Frontal on the snout / nose: 130 | Beak attachment point: 3 | |
| | Cheek: 2 | Beak: 1 | |
| | On the side of the snout / nose: 23 | Side of the Beak: 1 | |
| | No: 59 | No: 2 | |
| **Touches the stigma the deposited pollen?** | Yes (one observation) | No observation | No |
| **What is the spatial relationship between the stigma and pollen sacs to the animal?** | Not detectable: 561 | Not detectable: 22 | No interaction |
| | Over the cheek: 9 | Over the beak: 31 | |
| | Over the forehead:1 | Over the beak side: 1 | |
| | Side of the snout: 74 | Lateral to the beak: 2 | |
| | Lateral over the snout: 7 | Over the head and beak: 2 | |
| | Over the snout: 379 | Over the forehead: 3 | |
| **Position of the animal to the plant** | Frontally crouched to sideways, head tilted | Frontally crouched to sideways, head tilted | No interaction |
| **Interest of the animal** | Walks straight towards the plants | Move purposefully towards the plants. | Some walk towards the plants. Others walk past it. |
| **Does the animal cleans itself?** | Sand bathing: 1 | No | Paws over snout: 1 |
| | Tongue over snout: 3 | | |
| | Tongue + paws over snout: 2 | | |
| | Paws over snout: 5 | | |
| **Action** | Run through the video, drink nectar or sniff flowers. . | Drinks nectar or hops through video | Walk through the video or sniff flowers. Some nibbling on them. |
| **Damage to corolla, flower organs or whole plants** | No | No | No |
| **Paws as a tool?** | No | No | No |
