## Supplement 2 for "*Hyobanche hanekomii* (Orobanchaceae) is pollinated by non-flying mammals and birds"

**Table 2:** Result of the pairwise Wilcox test on the length of inflorescence visits for the flower visiting species on *H. hanekomii*.

| | ***E. edwardii* night** | ***E. edwardii* total** | ***E. edwardii* day** | **Muridae** |
|---|---|---|---|---|
| ***E. edwardii* total** | 0.980 | x | x | x |
| ***E. edwardii* day** | 0.980 | 0.980 | x | x |
| **Muridae** | 0.370 | 0.370 | 0.370 | x |
| ***N. famosa*** | 0.980 | 0.980 | 0.980 | 0.450 |

**Table 3:** Results of the pairwise Wilcox test on the length of the foraging bouts for the flower visiting species on *H. hanekomii*.

| | ***E. edwardii* night** | ***E. edwardii* total** | ***E. edwardii* day** | **Muridae** |
|---|---|---|---|---|
| ***E. edwardii* total** | 0.938 | x | x | x |
| ***E. edwardii* day** | 0.938 | 0.938 | x | x |
| **Muridae** | 0.036 | 0.036 | 0.038 | x |
| ***N. famosa*** | 0.938 | 0.938 | 0.938 | 0.120 |

**Table 4:** Result of the pairwise Wilcox test on the number of flower visits to drink nectar per foraging bout for the flower visiting species on *H. hanekomii*.

| | ***E. edwardii* night** | ***E. edwardii* total** | ***E. edwardii* day** | **Muridae** |
|---|---|---|---|---|
| ***E. edwardii* total** | 0.698 | x | x | x |
| ***E. edwardii* day** | 0.119 | 0.146 | x | x |
| **Muridae** | <0.001 | <0.001 | <0.001 | x |
| ***N. famosa*** | 0.006 | 0.006 | 0.027 | 0.005 |

**Table 5:** Result of the pairwise Wilcox test on the number of flower visits to for sniffing per foraging bout for the flower visiting species on *H. hanekomii*.

| | ***E. edwardii* night** | ***E. edwardii* total** | ***E. edwardii* day** | **Muridae** |
|---|---|---|---|---|
| ***E. edwardii* total** | 0.930 | x | x | x |
| ***E. edwardii* day** | 0.930 | 0.930 | x | x |
| **Muridae** | 0.930 | 0.930 | 0.930 | x |
| ***N. famosa*** | 0.690 | 0.690 | 0.690 | 0.930 |

**Table 6:** Result of the pairwise Wilcox test on the length of flower visits for drinking per foraging bout for the flower visiting species on *H. hanekomii*.

| | ***E. edwardii* night** | ***E. edwardii* total** | ***E. edwardii* day** |
|---|---|---|---|
| ***E. edwardii* total** | 0.568 | x | x |
| ***E. edwardii* day** | 0.041 | 0.063 | x |
| ***N. famosa*** | <0.001 | <0.001 | <0.001 |
